# ApoA-I dissociated from human HDL retains its acceptor properties in ABCA1-mediated cholesterol efflux from RAW 264.7 macrophages in coronary artery disease

**DOI:** 10.1101/2025.11.21.689699

**Authors:** Veronika B. Baserova, Mikhail A. Popov, Alexander D. Dergunov

## Abstract

**Background:** The significance of cholesterol efflux as a predictor of coronary artery disease (CAD) remains controversial. The primary route for intracellular cholesterol export via the ABCA1 transporter involves the acceptance of cholesterol by lipid-free apolipoprotein A-I and high-density lipoproteins (HDL). A separate estimate of the efficiencies of two reactions is thus required.

**Methods:** HDL from plasma of 63 control and 76 CAD male patients was obtained by PEG precipitation of apoB-containing lipoproteins and denatured by 4.25 M urea at the transition midpoint. We measured apoA-I dissociation concomitant with HDL denaturation, the expression of 65 preselected genes by real-time PCR, and the efficiency of ABCA1-mediated cholesterol efflux from RAW 264.7 macrophages to dissociated lipid-free apoA-I. The dissociation parameter *D* was calculated from the accumulation of apoA-I with preβ-mobility at HDL separation by agarose gel electrophoresis followed by immunodetection.

**Results:** The phospholipid:apoA-I and cholesterol: apoA-I ratios for CAD patients were consistently higher than those for control patients in a whole range of plasma HDL-cholesterol levels. ApoA-I partitioned 1.5-fold higher into the water phase for HDL from CAD patients relative to controls. For CAD patients, the dissociation parameter *D* was inversely correlated with phospholipid and cholesterol levels and the cholesterol:apoA-I ratio in HDL. For control patients, the *D* parameter was positively correlated with *CETP* and *ABCA1* gene expression levels. For CAD patients, the *D* parameter was negatively correlated with *CUBN* and *ALB* gene expression levels that may be associated with the increased catabolism of lipid-free apoA-I. The *V*_m_ and *K*_m_ values showed that apoA-I functionality in ABCA1-mediated cholesterol efflux from RAW 264.7 macrophages to lipid-free apoA-I generated at urea-induced HDL denaturation was similar for HDL from both groups.

**Conclusions:** The enrichment of HDL with cholesterol, with the concomitant increase of competition between apoA-I and cholesterol for the binding to phospholipid molecules adjacent to apoA-I, could be involved in the increased apolipoprotein dissociation from HDL in CAD. The obtained estimates for the ABCA1-mediated cholesterol efflux to lipid-free apoA-I may be the prerequisite for the detailed study of cholesterol efflux kinetics with two apoA-I forms that could allow to be described them as CAD predictors.

## 1. Introduction

The human plasma high-density lipoproteins (HDL) possess an atheroprotective effect through the reverse cholesterol transport from arterial-wall macrophages to the liver [1] that begins with cholesterol efflux via active cholesterol transport by ABCA1 and ABCG1 transporters, facilitated diffusion with SR-B1, and passive diffusion [2]. The apoA-I is a major HDL apolipoprotein that exists in lipid-bound and lipid-free forms; the latter is a primary cholesterol acceptor in the ABCA1-mediated efflux [3]. However, small HDL particles are efficient cholesterol acceptors as well [4]. The HDL heterogeneity by density and charge results in the existence of light HDL_2_ and dense HDL_3_ and particles with preβ-mobility and α-mobility, respectively. Lipid-free apoA-I is found only in the preβ-band [5], which also includes nascent HDL, while mature spherical HDL is localized in the α-band. The nascent HDL, as the initial product of cholesterol and phospholipid efflux by ABCA1, further accepts these lipids with the formation of disc-shaped particles. The latter are the most efficient substrate for lecithin:cholesterol acyltransferase (LCAT). The enzyme converts free cholesterol to cholesteryl ester that is thought to prevent the reabsorption of effluxed cholesterol. The subsequent steps of reverse cholesterol transport include the transfer of cholesteryl ester from HDL to larger very low-density and low-density lipoproteins (LDL) by cholesteryl ester transfer protein (CETP), the SR-B1-mediated capture of cholesteryl ester, and the uptake of a whole LDL particle by the LDL receptor in the liver [2]. The remodeling of spherical HDL by CETP and lipases results in a spherical-to-discoidal transformation with a concomitant transition between lipid-free and lipid-bound apoA-I [6]. Of note, the dissociated apoA-I [7; 8] and HDL-apoA-I exchange [9] may be potentially involved in cholesterol efflux.

A strong inverse relationship between HDL cholesterol efflux capacity (CEC) and coronary artery disease (CAD) signifies HDL functionality more than HDL-cholesterol level in the HDL atheroprotective effect [10]. HDL functionality includes HDL particle number and heterogeneity and particle ability to accept effluxed cholesterol. The significance of ABCA1-mediated efflux and HDL functionality in CAD are inconsistent [4; 11–16]. CEC from macrophages had a strong inverse association with morphological and prognostic markers of angiographic CAD [17]. The higher concentration of preβ_1_ particles was responsible for the lower preβ_1_ concentration-normalized ABCA1-dependent efflux capacity in CAD [18; 19]. ABCA1-dependent cholesterol efflux was positively correlated with the levels of small lipid-poor preβ_1_ particles [18]. At variance with this, HDL efflux capacity was inversely associated with small HDL particle concentration in older adults [20]. Furthermore, the association between HDL particle size and composition and CAD risk is not conclusive, and the inclusion and magnitude of lipid-free apoA-I and HDL particles in individual steps of cholesterol efflux remain to be implicitly solved.

We describe now the relations between apoA-I dissociation, HDL functionality and the expression of selected genes controlling HDL metabolism and atherogenesis in control and CAD patients.

## 2. Materials and Methods

Patient selection with widely varied HDL-C levels (63 control patients and 76 patients with CAD confirmed by coronary angiography) and laboratory tests were described previously [21]. The crude HDL preparations (pHDL) were prepared by the precipitation of apoB-containing lipoproteins with PEG 7000–9000 [7]. The concentrations of choline-containing phospholipids and total cholesterol in HDL preparations were measured by enzyme methods using Sentinel (Italy) and HUMAN (Germany) kits, respectively, and apoA-I was measured by immunonephelometry with AU 480 (Beckman Coulter, USA).

ApoA-I content in preβ- and α-fractions of pHDL was measured by agarose gel electrophoresis followed by immunodetection of apoA-I [7]. PeakFit version 4.12 (SeaSolve Software Inc.) and ImageJ software were used to determine band intensities with baseline subtraction at the conditions of linear response of each band intensity to the amount of loaded sample.

Urea-induced pHDL denaturation after incubation for 6 h at 25 °C was followed by the accumulation of lipid-free preβ-apoA-I detected in the immunoreplica of agarose gel. ApoA-I dissociation was analyzed from the fractions of preβ-signal detected after the treatment with 0 M, 4.25 M, and 7.5 M urea [7]. The degree of apolipoprotein dissociation at 4.25 M urea as a mid-transition zone that is accomplished at 7.5 M urea was characterized by dissociation parameter *D* (Equation 1):

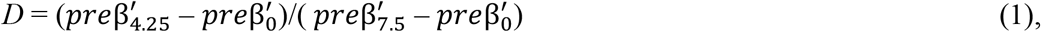

Where 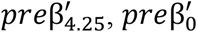, and 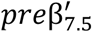 are preβ fractions with 4.25 M, 0 M, and 7.5 M urea, respectively. The dependence of urea-induced apoA-I release on phospholipid level in pHDL preparations was characterized by the partition coefficient *K* between aqueous and lipid phases adjusted for water molarity (Equation 2):

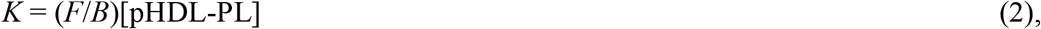

where *F* and *B* are the numbers of molecules of lipid-free apoA-I in water and bound to pHDL phospholipids, respectively [22]. The *F*/*B* ratio was calculated from the *D* parameter measured at 4.25 M urea (Equation 3):

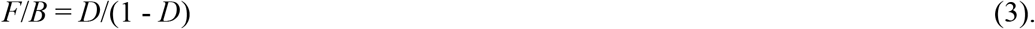

Cholesterol efflux from murine macrophages RAW 264.7 to pHDL was measured with a fluorescent probe BODIPY-cholesterol. The basal and cAMP-stimulated effluxes were measured, and ABCA1-mediated efflux was calculated as a difference between stimulated and basal effluxes and expressed as a percent of effluxed cholesterol [7].

To separate the contributions of both preβ-HDL and mature α-HDL in ABCA1-mediated efflux, a simple kinetic theorem for two-substrate reaction, when substrates react with a single enzyme without the formation of any ternary complex [23], was applied for a total value of ABCA1-mediated efflux with pHDL as cholesterol acceptor. The total initial rate of reaction *v*_*tot*_ when preβ-HDL and α-HDL act as a competitive inhibitor of the other for the binding to ABCA1 transporter is described by the Equation 4:

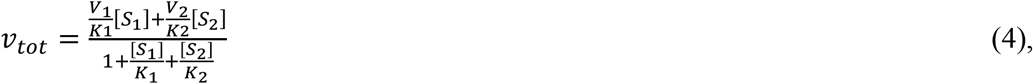

where *V*_i_, *K*_i_, and [*S*_i_] are the maximal velocities, Michaelis constants and substrate concentrations for preβ-HDL and α-HDL as a first and second substrates, respectively, with the relation *V*_1_ > *V*_2_. Both [*S*_1_] and [*S*_2_] are apoA-I concentrations in preβ-band and α-band. The treatment of Equation 4 in the double reciprocal plots for a first substrate at a constant [*S*_2_]/[*S*_1_] ratio *R* results in the Equation 5:

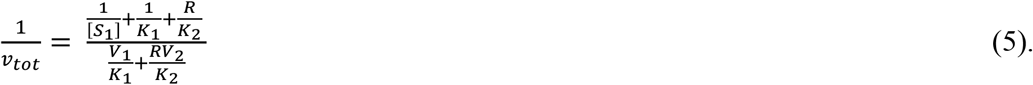

For series of lines in double reciprocal plots for a first substrate with various *R* values the common interception point does not depend on *R* and abscissa and ordinate values are determined by Equations 6 and 7, respectively:

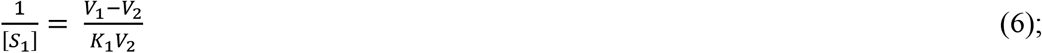

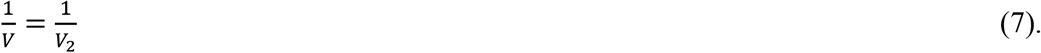

A common maximal velocity *V*_m_ at the constant *R* is determined by the Equation 8:

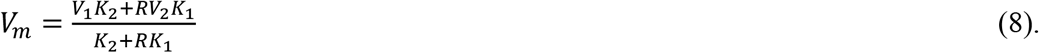

In turn, the *K*_2_ value is derived from Equation 8 (Equation 9):

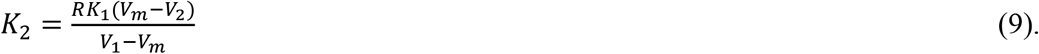

*V*m varies between low *V*_2_ and high *V*_1_ borders. Generally, *V*m non-linearly depends on *R*. The linear plots according to Equation 5 were constructed for fully denatured and intact pHDL sets and the data were subsequently treated with the equations 6-9 to derive kinetic parameters. The *V*_1_ and *K*_1_ values were measured without *S*_2_ interference with pHDL fully denatured with 7.5 urea. Before mixing, predenatured HDL were diluted 1:100 to lower urea concentration. The residual urea concentration not exceeding 75 mM did not influence efflux

The expression of 65 genes involved in HDL metabolism and atherogenesis measured by us earlier by real-time PCR in peripheral blood mononuclear cells [21; 24] was compared to apoA-I dissociation from pHDL in the present study.

Statistica 13 software was used for statistical analysis. The associations between variables were analyzed by the Pearson correlation coefficient. The statistical significance limit was accepted as *p* < 0.05. Nonlinear curve fitting was done with OriginPro 9.0.0 SR1. The significance of the difference between different dataset fits was checked with the F-statistic.

## 3. Results

### 3.1. Relations between ApoA-I Dissociation, HDL Composition, and Gene Expression

The dissociation of apoA-I from pHDL induced by urea treatment and measured by the *D* parameter significantly decreased with the increase of absolute and apoA-I-normalized cholesterol levels in pHDL from CAD patients (Table 1). The apoA-I dissociation changed quite analogously with the increase of absolute levels of choline-containing phospholipid (PL). The fit of dissociation data, passed through zero, to the linear model of apoA-I distribution between lipid-bound (B) and lipid-free (F) states (Equation 2) results in the values of the partition coefficient between aqueous and lipid phases adjusted for water molarity *K* = 0.239 ± 0.012 (R^2^ = 0.874) for control pHDL and *K* = 0.368 ± 0.020 (R^2^ = 0.822) for pHDL from CAD patients (Fig. 1). Thus, a 1.54-fold prevalence of apoA-I distribution between water and lipid phases exists in CAD relative to controls (*p* = 0.000) with the value of the difference in the standard free energy 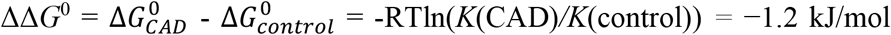, the difference value being positive for apolipoprotein distribution between lipid and water phases. ApoA-I in HDL from CAD patients possesses higher free energy and, thus, exhibits a higher dissociation. Of note, the increased PL:apoA-I and Chol:apoA-I ratios for pHDL from CAD patients relative to controls were measured throughout the whole range of HDL-C concentration in both patient groups (data not shown).

**Table 1.**
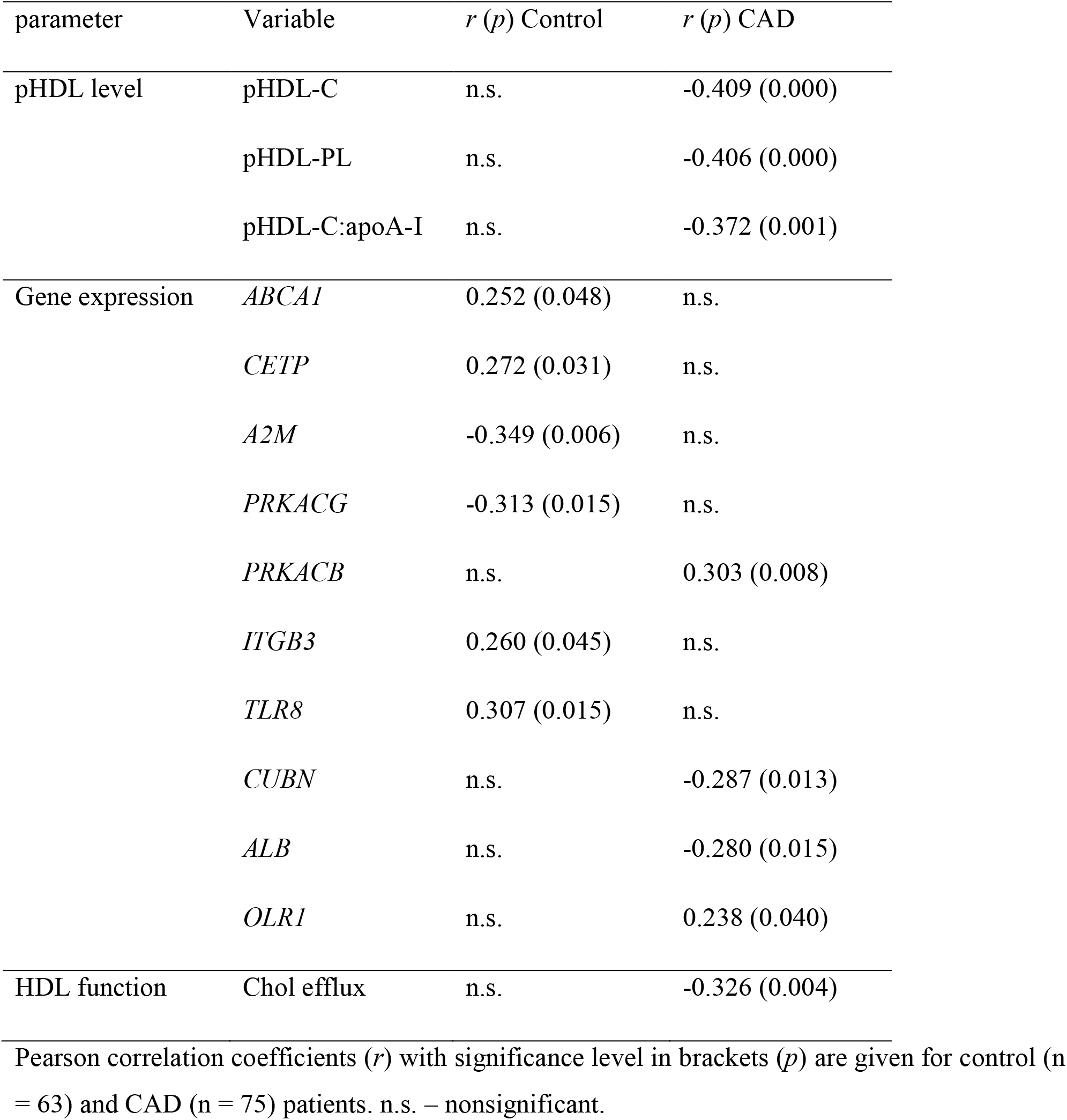
Relation of dissociation parameter *D* with pHDL lipids, gene expression, and ABCA1-mediated cholesterol efflux.

**Figure 1.**
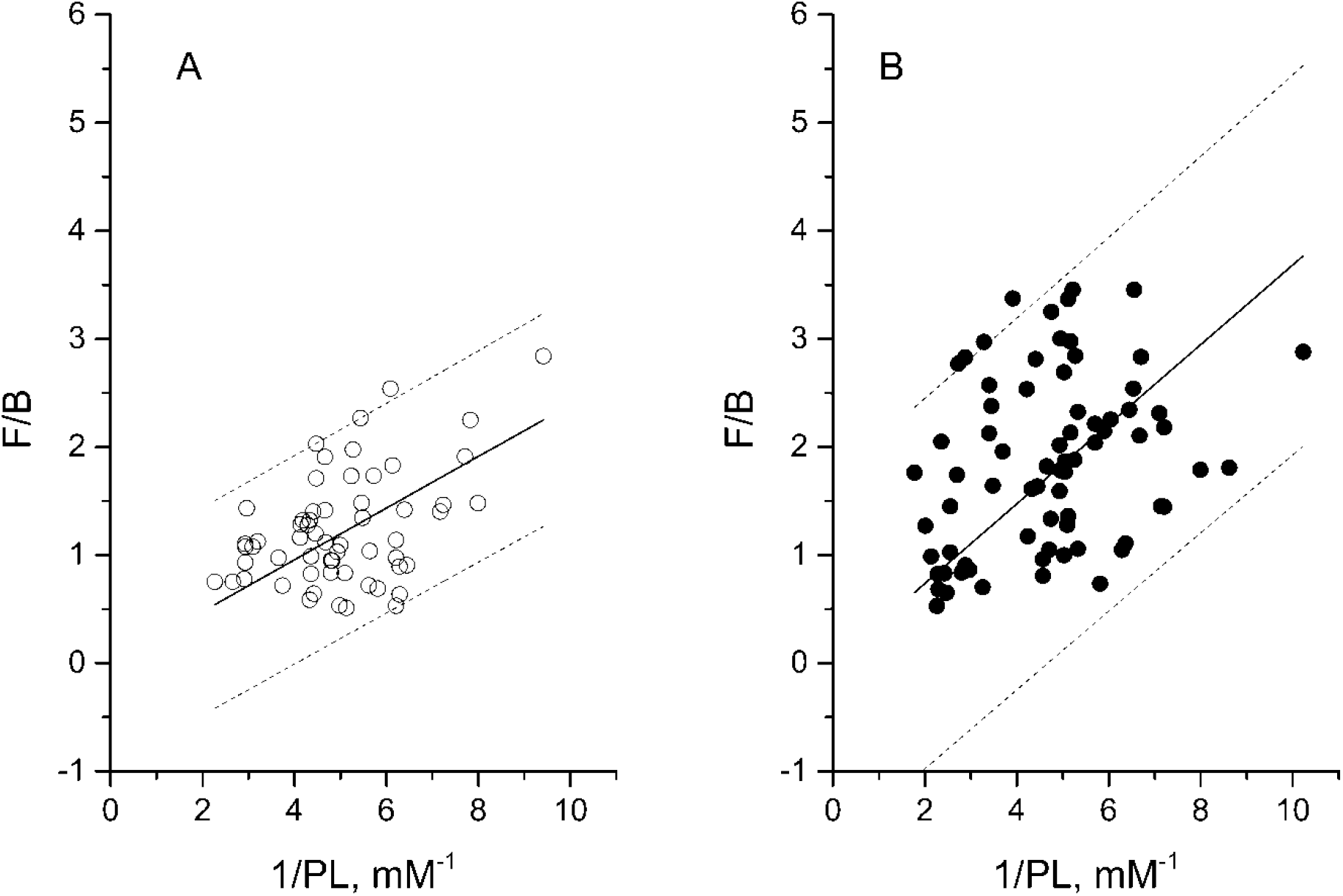
ApoA-I distribution between water and lipid phases of pHDL predenatured by 4.25 M urea. (A) pHDL from control patients (n = 59), (B) pHDL from CAD patients (n = 73). The 95% prediction bands are given as thin dashed lines. The Y-intercepts passed through zero according to the distribution model. Slopes were 0.239 ± 0.012 (R^2^ = 0.874) for control and 0.368 ± 0.020 (R^2^ = 0.822) for CAD. Slope differences were significant with *p* = 0.000.

To reveal the associations between apoA-I dissociation and lipoprotein metabolism, the expression of 65 genes in peripheral blood mononuclear cells, sensitive to HDL metabolism and atherogenesis, was analyzed. The significant correlations between *D* and gene expression are included in Table 1. For control patients, the positive correlation of the *D* parameter with the *CETP* expression level existed. The positive correlation of the *D* parameter with the *ABCA1* expression level may be associated with the decrease of lipid-free apoA-I with the increase of ABCA1 activity and the accompanying increase of dissociation of lipid-bound apolipoprotein. The *A2M* and *PRKACG* expression levels were negatively correlated with the *D* parameter. Remarkably, the increased expression of the *A2M* gene has been reported to be the marker of atherosclerosis progression [25]. This suggestion contrasts with the positive correlation of the dissociation parameter with the expression of the *ITGB3* and *TLR8* genes which possess the proatherogenic effect in CAD [24]. For patients with CAD, the apoA-I dissociation from pHDL is not associated with *CETP* expression level. However, the negative correlations of the *D* parameter with *CUBN* and *ALB* expression levels existed.

### 3.2. Relations between ApoA-I and Cholesterol Efflux

First, the efflux measurements were done with intact pHDL as cholesterol acceptors. For controls, there was no significant correlation between apoA-I dissociation and ABCA1-mediated cholesterol efflux. In CAD, efflux was less efficient relative to controls, and dissociation parameter *D* was negatively correlated with efflux (Table 1).

Second, due to the involvement of lipid-free and lipid-bound apoA-I in cholesterol transport by the ABCA1 transporter, the decrease of efflux efficiency in CAD may be associated with the less efficient efflux with each or both acceptors. To separate individual contributions, the functionality of lipid-free apoA-I was studied. To induce apoA-I dissociation from the HDL lipid phase to the water phase, pHDL from control and CAD patients were predenatured for 6 h at 25°C by 4.25 M urea at the middle of the HDL denaturation transition. The ABCA1-mediated efflux for 2 h to serially diluted pHDL was then measured, and the data were fitted to Michaelis-Menten kinetics (Fig. 2). The kinetic parameters are given in Table 2. The concentration of lipid-free preβ-apoA-I dissociated from mature α-HDL was calculated as a product of dissociation parameter *D* and apoA-I level in the α-band estimated by agarose gel electrophoresis and apolipoprotein immunodetection (*D* × α-apoA-I). The dependencies were well fitted to the equation for one substrate reaction and did not differ by the F-test for control and CAD. The *V*_m_, *K*_m_, and *V*_m_/*K*_m_ kinetic parameters did not differ also for pHDL from control and CAD patients by Student’s *t*-test. Of note, the kinetic parameters, checked by the F-statistic, did not change significantly with pHDL predenatured with 7.5 M urea (Table 2) at the completion of the denaturation transition for pHDL from control or CAD patients, which assumes no contribution of HDL remaining at the transition midpoint into the cholesterol efflux. Thus, the lipid-free apoA-I seems to be equally efficient as an acceptor of cholesterol effluxed by ABCA1 in CAD relative to control samples.

**Table 2.** Kinetics of ABCA1-mediated efflux to pHDL predenatured by urea.

| patient | Urea, M | n | V<sub>m</sub>, %/2h | K<sub>m</sub>, µg/ml | V<sub>m</sub>/K<sub>m</sub> |
| --- | --- | --- | --- | --- | --- |
| Control | 4.25 | 29 | 16.6 ± 3.5 | 0.38 ± 0.15 | 44.2 ± 19.7 |
|  | 4.25 | 5 | 16.8 ± 3.5 | 0.69 ± 0.27 | 24.4 ± 10.9 |
|  | 7.50 | 5 | 15.0 ± 1.9 | 0.74 ± 0.22 | 20.3 ± 6.6 |
| CAD | 4.25 | 31 | 16.5 ± 3.9 | 0.47 ± 0.20 | 35.0 ± 16.9 |
|  | 4.25 | 4 | 18.9 ± 1.6 | 0.61 ± 0.11 | 30.8 ± 6.0 |
|  | 7.50 | 4 | 17.7 ± 1.0 | 0.62 ± 0.08 | 28.3 ± 4.1 |

Mean ± SE values are given. n, patient number.

**Figure 2.**
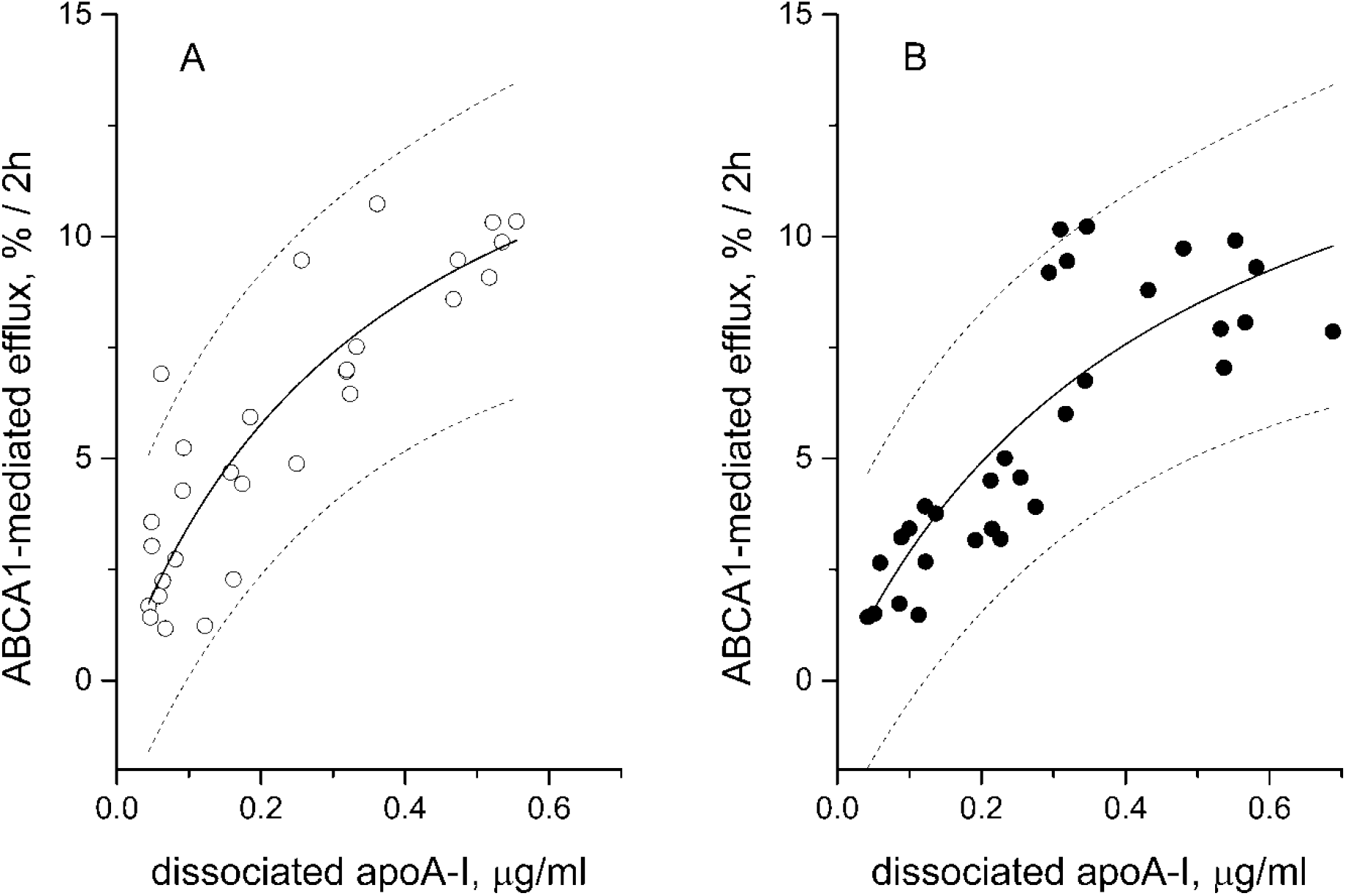
The kinetics of ABCA1-mediated efflux to pHDL predenatured by 4.25 M urea. (A) pHDL from control patients (n = 29), (B) pHDL from CAD patients (n = 31). The 95% prediction bands are given as thin dashed lines.

We used this approach to separate the contributions of lipid-free and HDL-bound apoA-I as first and second substrates, respectively, to the kinetics of ABCA1-mediated efflux based on the simple kinetic theorem of two substrates competing for one enzyme (Equation 4). The initial rates of efflux to intact and fully denatured pHDL from CAD patients with low HDL-C were treated as a function of preβ-apoA-I concentration [*S*_1_] in double reciprocal plots (Fig. 3). The major prerequisite for a plot linearity is a constant [*S*_2_]/[*S*_1_] ratio *R* (Equation 5). The efflux was measured separately with two intact pHDL with similar R values and five denatured pHDL, and the data were concatenated to increase the statistical power. The measurements with denatured pHDL were necessary to derive *V*_1_ and *K*_1_ values without *S*_2_ interference. The *V*_2_ and *K*_2_ values for intact pHDL with *R* value 13.0 were derived with Equations 6-9. The complete set of mean values of kinetic parameters included *V*_1_ = 17.5 %/2h, *K*_1_ = 0.62 µg/ml, *V*_2_ = 5.3 %/2h, and *K*_2_ = 2.13 µg/ml. Thus, efflux efficiency, expressed as the *V*_m_/*K*_m_ ratio, for lipid-free apoA-I was 11.4-fold higher than that for HDL-bound apoA-I in HDL from CAD patients with low HDL-C level.

**Figure 3.**
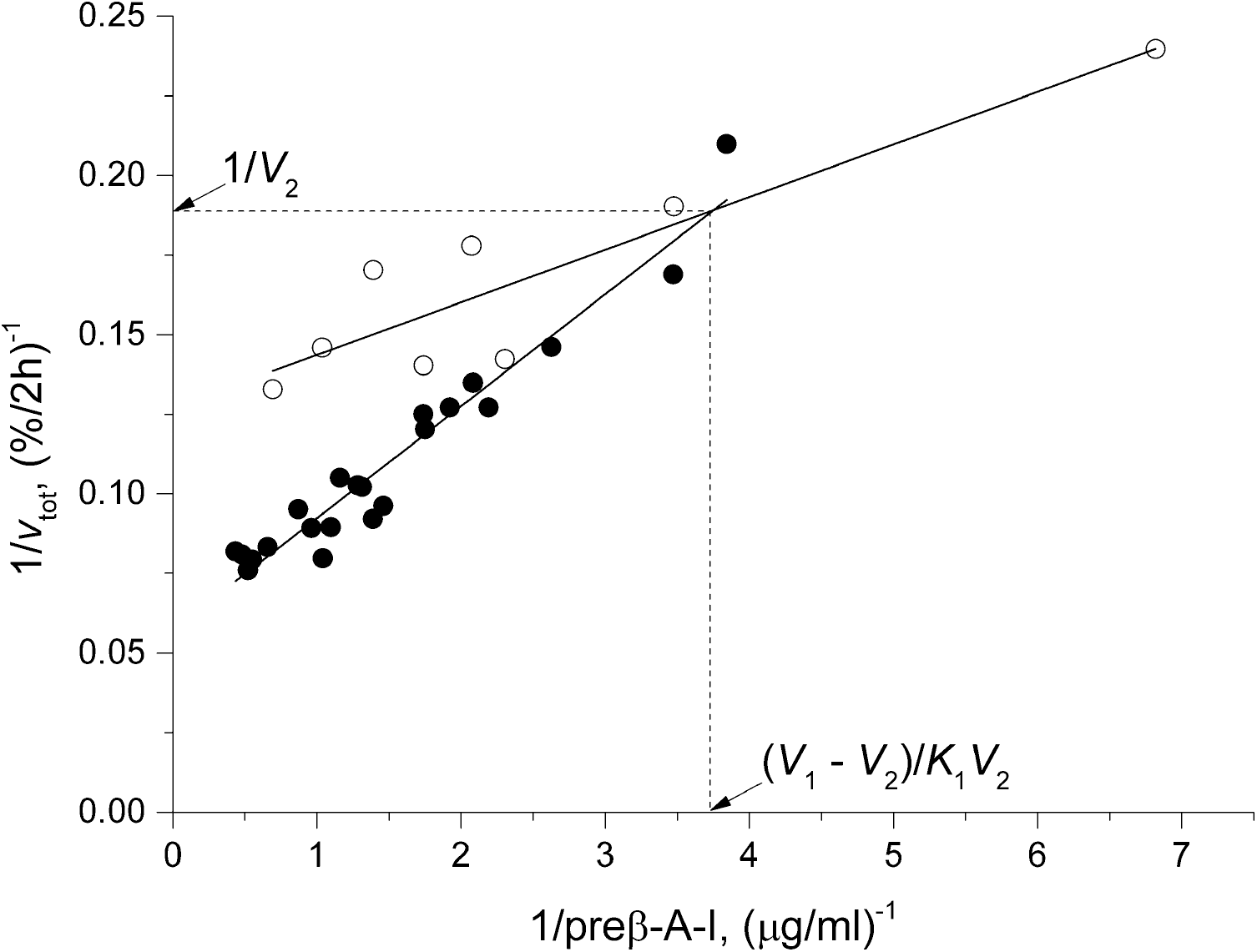
The dependency of the total initial rates of ABCA1-mediated cholesterol efflux *v*_tot_ on *S*_1_ concentration in double reciprocal plots for intact and denatured pHDL from the CAD group with low plasma HDL-C levels. The data were fitted to a linear function 1/*v*_tot_ = *intercept* + *slope* × (1/[*S*_1_]). The efflux was measured with five individual pHDL pre-denatured with 7.5 M urea (filled symbols), and the data were concatenated for fit: the *intercept* was 0.0573 ± 0.003 (*p* = 0.000), the *slope* was 0.035 ± 0.002 (*p* = 0.000), and *R*^2^ = 0.94. For two intact pHDL with a mean value of *R* = 13.0 (open symbols), the *intercept* was 0.127 ± 0.009 (*p* = 0.000), the *slope* was 0.017 ± 0.003 (p = 0.002), and *R*^2^ = 0.83. The ordinate and abscissa of the intersection point were used to calculate the kinetic parameters with *S*_2_.

## 4. Discussion

There are three major findings in the present study revealed at the comparison of compositional and functional properties of HDL from CAD and control patients. First, for HDL from CAD patients, apoA-I partitioned more into the water phase during urea-induced HDL denaturation, and distribution parameter *D* was inversely associated with the absolute level of choline-containing phospholipid and the absolute and apoA-I-normalized cholesterol levels in pHDL, opposite to the lack of any association for HDL from control patients. The increased denaturation of HDL from the patients with acute coronary syndrome has been described [26]. Importantly, HDL particles from CAD patients are more enriched with PL and Chol normalized by apoA-I level throughout the whole range of HDL-C. The enrichment of HDL with cholesterol, with the concomitant increase of competition between apoA-I and cholesterol for the binding to phospholipid molecules adjacent to apoA-I, is suggested to be involved in the increased apolipoprotein dissociation from HDL in CAD. Cholesterol molecules have been shown to be excluded partially from boundary lipid in reconstituted HDL [27]. The free cholesterol molecules may increasingly accumulate in the vicinity of apoA-I molecules in cholesterol-overloaded pHDL from CAD patients with the additional perturbation of PL structural order and dynamics. What could be a driving force for additional cholesterol accumulation in HDL from CAD patients compared to control patients? We measured earlier the upregulated expression of *CETP, LPL*, and *PLTP* genes and down-regulated expression of *LCAT* gene in CAD patients [24]. The differential expression of these genes is suggested to be involved in lipid enrichment of HDL in CAD. Indeed, the enrichment of HDL with phospholipid and cholesterol molecules in CAD could occur due to a decreased phospholipid hydrolysis by LCAT which possesses phospholipase activity with a concomitant lowering of cholesteryl ester generation [28]. The entropically favourable processes of both apoA-I exchange between discoidal HDL-bound and lipid-free pools [29] and the lipid transfer for nanodisks consisting of apoA-I and DMPC [30] with the increased activation entropy and the decrease in the standard entropy have been suggested. The exact contribution of entropy and enthalpy into apoA-I increased dissociation from mature spherical HDL in CAD remains to be determined. The different exposure of the hydrophobic amino acid residues of apoA-I to the aqueous phase in the activated state at apolipoprotein denaturation [22] may be invoved. Interestingly, cholesterol in discoidal HDL increased the number of apoA-I tryptophan residues accessible to the aqueous phase, but decreased their mean degree of hydration [31].

Second, for HDL from control patients, the *D* parameter was positively correlated with the *CETP* expression. CETP-induced apoA-I dissociation from HDL from healthy normolipidemic subjects has been described by Barter et al. [32;33]. Also, triglyceride enrichment of HDL core, that augments apolipoprotein dissociation from the surface [34] may underlie the positive association between the *D* parameter and *CETP* expression. Of note, the increased apoA-I dissociation seems to possess a proatherogenic effect in control patients due to the positive associations between the *D* parameter and *ITGB3* and *TLR8* gene expression. For HDL from CAD patients, the *D* parameter was not associated with *CETP* expression. However, the observed negative associations of the *D* parameter with *CUBN* and *ALB* gene expression may be associated with the increased catabolism of lipid-free apoA-I and the accompanying decrease of the preβ fraction in CAD. Indeed, CUBN as an endocytic receptor is involved in endocytosis of apoA-I and albumin and maintenance of their blood levels [35]. Interestingly, the reciprocal associations existed between the dissociation parameter and the expression levels of two genes, i.e. negative for *PRKACG* for control patients and positive for *PRKACB* for CAD patients. The expression levels of these genes contributed reciprocally to HDL-C variability in CAD patients [24].

Third, lipid-free/lipid-poor apoA-I that may originate from the HDL-bound state both for CAD and control HDL, seems to be equally efficient in ABCA1-mediated efflux. The obtained *V*_m_ and *K*_m_ values are lower than the analogous mean values (22.3% and 0.64 µg/ml) for ABCA1-mediated efflux to lipid-free apoA-I for 6 h [4]. Of note, only 40-60% of apoA-I molecules are dissociated in water upon pHDL denaturation by 4.25 M urea (compared to 10-20% of preβ-apoA-I in intact HDL), and *V*_m_ could not reach the maximal limiting value for lipid-free apoA-I due to the lower *V*_m_ value for remaining lipid-bound apoA-I. However, similar efflux kinetics at two zones of denaturation transition largely exclude this possibility. Of note, the structure of residual HDL with lipid-bound apoA-I, still existing at the transition midpoint, is perturbed by urea, which excludes the possibility of accepting cholesterol by residual HDL.

The decreased ABCA1-mediated efflux efficiency of intact HDL as a mixture of lipid-poor and lipid-bound apoA-I in CAD has been described [18;19]. Despite the attempts to normalize efflux to preβ_1_-levels as an indicator of lipid-poor apolipoprotein, the conventional CEC measurements are performed with HDL as a mixture of lipid-poor and lipid-bound apoA-I without the separation of individual efflux reactions with different acceptors [18;19;36]. However, the overall reaction velocity for a two-substrate reaction with a single enzyme depends on the concentrations of both substrates depicted by *R* values (Equation 8), and the numerical simulation of this dependency revealed a non-linear increase of the *V*_m_ value with the decrease of the *R* value. The effective value of the maximal reaction velocity *V*_m_ 7.9 %/2h for intact HDL from CAD patients with low HDL-C level and with the mean *R* value 13.0 lies between *V*_1_ (17.5 %/2h) and *V*_2_ (5.3 %/2h) limits. The exact nature of the dependency of *V*_m_ on *R* for control and CAD patients with different HDL-C levels is a matter of ongoing experiments based on the application of kinetics of lipid-free and HDL-bound apoA-I competing with each other for the ABCA1 transporter. We suggest that this dependency may underlie the diminished cholesterol efflux in CAD.

## 5. Conclusions

For control patients, the dissociation of apoA-I in the water phase at urea-induced HDL denaturation is associated with *ABCA1* and *CETP* gene expression. For CAD patients, apoA-I partitions more into the water phase due to the higher free energy of apolipoprotein in the lipid phase with the negative correlations of the dissociation parameter with normalized cholesterol levels in HDL particles. The total initial rate of ABCA1-mediated cholesterol efflux from RAW 264.7 macrophages, when preβ-HDL and α-HDL act as competitive inhibitors of each other for the binding to ABCA1 transporter, was measured with intact HDL and pre-denatured HDL as a source of lipid-free apoA-I. The retained CEC of lipid-free/lipid-poor apoA-I in CAD may be masked by the higher competition of α-HDL, which possesses lower CEC, with preβ-HDL for ABCA1 binding. This new approach may be used in the analysis of lipid-free and HDL-bound apoA-I as independent predictors of CAD.

## Contributions

A.D. Dergunov – concept of the study; writing text of the paper; supervision of the study; M.A. Popov – patients; V.B. Baserova – conducting experiments; preparation of figures and tables. All authors read and approved final version of the manuscript.

## Ethics approval and consent to participate

All procedures performed in studies involving human participants were in accordance with the ethical standards of the institutional and/or national research committee and with the 1964 Helsinki declaration and its later amendments or comparable ethical standards. The project was approved by ethics committee of the M.F. Vladimirsky Moscow Regional Research and Clinical Institute MONIKI (protocol no. 12479/2019, February 17, 2019). All involved patients provided voluntary informed consent to participate in the study.

## Declaration of generative AI and AI-assisted technologies in the writing process

During the preparation of this work the authors did not use AI in the writing process.

